# Explicitly nonlinear fMRI networks reveal hidden trajectories of infant brain development

**DOI:** 10.64898/2026.04.07.716703

**Authors:** Spencer Kinsey, Geethanjali Nagaboina, Prerana Bajracharya, Masoud Seraji, Zening Fu, Vince D. Calhoun, Sarah Shultz, Armin Iraji

## Abstract

Nonlinearity is a hallmark of brain complexity at multiple scales. However, existing functional magnetic resonance imaging (fMRI) functional connectivity studies typically utilize linear methods. Therefore, links between nonlinear fMRI connectivity patterns and the development of the human brain during critical periods such as infancy remain unclear. To address this gap in knowledge, we developed a data-driven approach to capture brain intrinsic connectivity networks from explicitly nonlinear resting-state fMRI connectivity and profiled their developmental associations in a cohort of typically developing human infants. We identified neurobiologically structured nonlinear fMRI connectivity patterns during early postnatal life, indicating that macroscopic brain ensembles systematically participate in nonlinear relationships at birth. Furthermore, we found that linear and explicitly nonlinear network counterparts are linked to partially overlapping but complementary developmental profiles during this period of rapid brain maturation, with the explicitly nonlinear approach unveiling insights into the development of networks that have been associated with sensorimotor capacities, default mode processes, executive functioning, language production, and stimulus saliency. Our study marks the first comprehensive developmental investigation of whole-brain nonlinear fMRI networks in human infants and deepens contemporary perspectives on neuroimaging data discovery by emphasizing the informational richness of nonlinear relationships at fMRI scales of observation.

## Introduction

How large-scale functional brain organization evolves during early human development remains a fundamental open question in neuroscience. Infancy demarcates a critical period of rapid human brain maturation characterized by myelination and synaptogenesis (Gilmore et al., 2018; Natu et al., 2021) which may serve as a reservoir of early biomarkers for neurodevelopmental and/or psychiatric risk. Given the brain’s plasticity during infancy, altered developmental trajectories during this period can impact behavior throughout childhood, adolescence, and adulthood. However, early functional brain organization remains incompletely understood.

Functional magnetic resonance imaging (fMRI) functional connectivity (FC) research can help address these gaps by allowing researchers to non-invasively characterize coherent networks of neurally relevant (Logothetis et al., 2001) blood-oxygenation-level-dependent (BOLD) signal fluctuations in unprecedented comprehensive fashion. FC research has been identified as an important emerging direction in the study of human brain development (Desrosiers et al., 2024; Power et al., 2010; Yin et al., 2025) with implications for neurodevelopmental, psychiatric, and clinical risk screening and assessment. However, most FC analyses in fMRI continue to rely on methods that privilege linear measures such as Pearson correlation (Friston, 2011) with the implicit assumption of linear, Gaussian relationships. While these approaches have yielded important insights, this assumption constrains the types of statistical dependencies that can be detected and may therefore overlook features of functional organization that deviate from linear structure.

Several factors contribute to the predominance of linear FC methods: nonlinear methods can be computationally demanding and less straightforward to implement at scale (Jin et al., 2025), the statistical and spatiotemporal characteristics of fMRI data may limit the added value of nonlinear estimators according to some criteria, as suggested by Hlinka et al. (2011) and Raffaelli et al. (2024), and linear metrics generally offer more transparent interpretability relative to nonlinear approaches (Friston, 2001). However, several studies have shown that potentially informative nonlinear relationships in fMRI data are under-recognized due to methodological choices that prioritize linear approaches (Iraji et al., 2023a; Kinsey et al., 2024; Motlaghian et al., 2022). For example, networks can be reliably inferred from voxel-scale nonlinear fMRI FC patterns even after removing linear FC information, and such networks exhibit enhanced statistical sensitivity to clinical conditions such as schizophrenia relative to their linear counterparts (Iraji et al., 2023a; Kinsey et al., 2024). These observations suggest that applications of nonlinear estimation methods at the macroscale should be assessed according to their potential to provide insights into the underlying neurofunctional organization of the brain, as even subtle nonlinear fMRI FC patterns might reflect biophysical properties of neural populations that support phenomena such as harmonic generation and cross-frequency coupling (Wang et al., 2025). Given that neurophysiological properties may change rapidly during critical periods (Reh et al., 2020), the assumption of linear, Gaussian FC may be particularly limiting in studies of infant brain development.

The relatively few nonlinear fMRI FC studies that exist have focused on adults and have typically analyzed relationships between time series of predefined regions of interest (ROIs) rather than voxel time series (Hlinka et al., 2011; Jin et al., 2025; Raffaelli et al., 2024). However, the spatiotemporal averaging involved in ROI time series extraction inherently emphasizes linearity and might obscure informative nonlinear dependencies. Moreover, the use of predefined (rather than data-driven) nodes risks averaging across the signals of brain areas that exhibit distinct types of functionality, potentially further de-emphasizing informative nonlinear relationships. Finally, the practice of defining networks using *a priori* atlases does not address the fundamental need to infer brain systems that optimally reflect the data’s FC structure and more precisely capture meaningful participant-level variation (Calhoun et al., 2009; Gordon et al., 2017; Korhonen et al., 2021).

In this study, we address these limitations by proposing a multivariate data-driven method to learn networks from voxel-wise explicitly nonlinear (ENL) FC patterns, allowing us to profile previously hidden developmental trajectories of nonlinear fMRI synchronization in the infant brain (Fig. 1).

**Fig. 1.**
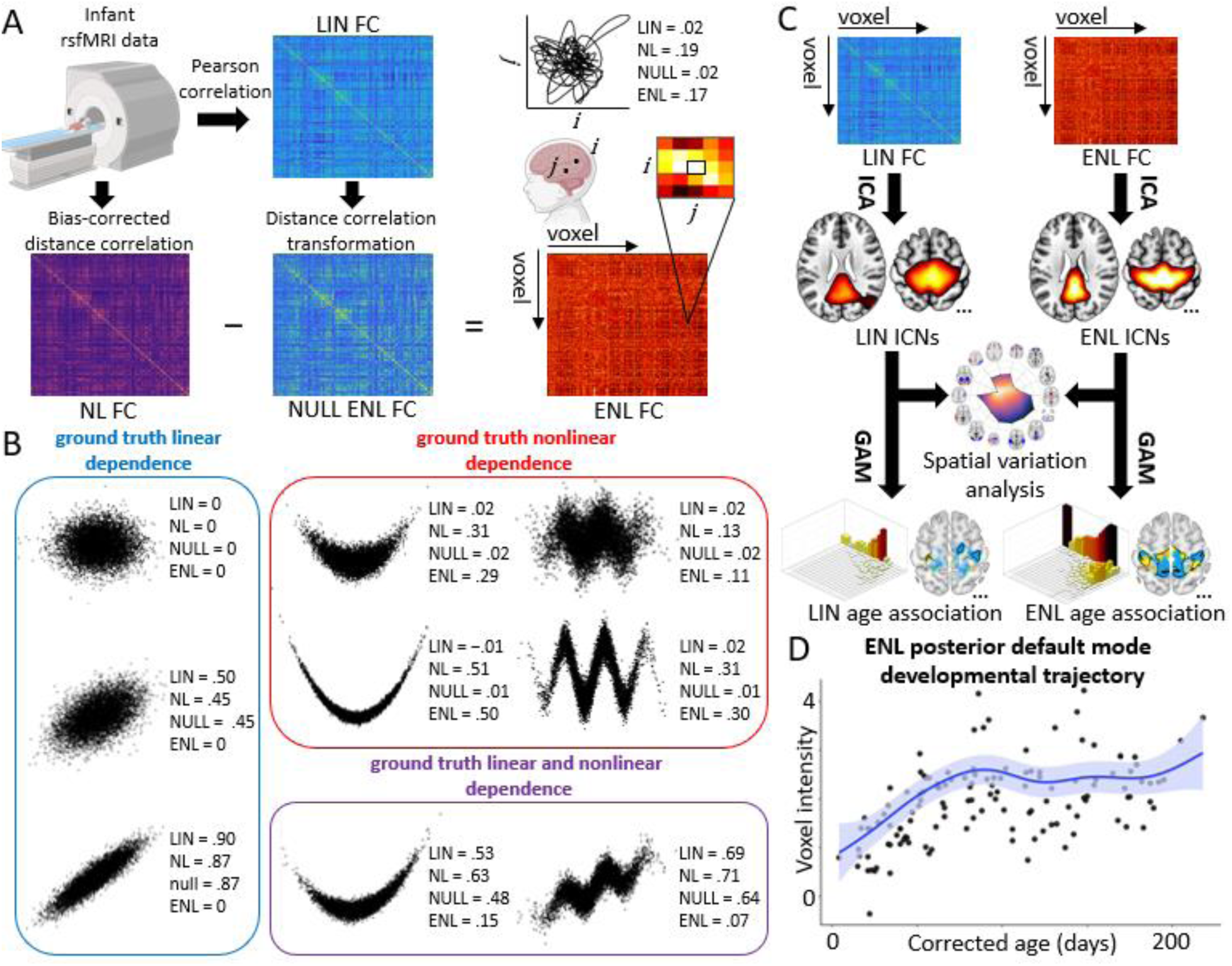
Methodology for extracting explicitly nonlinear (ENL) intrinsic connectivity networks (ICNs) from resting-state fMRI (rsfMRI) data and the subsequent identification of developmental profiles. (A) Voxel-level rsfMRI data was first transformed into linear (LIN) functional connectivity (FC) via Pearson correlation. LIN FC was then transformed into the distance correlation metric space under the assumption of bivariate Gaussianity, yielding the expected whole-brain distance correlation matrix if only linear relationships were present (i.e., the multivariate NULL ENL FC matrix). To capture nonlinear (NL) voxel-by-voxel FC, the bias-corrected distance correlation was computed. Finally, NULL ENL FC was subtracted from NL FC, yielding ENL FC at the whole-brain level. The joint activity trajectory of two voxels is depicted as an example of high nonlinearity in a relationship which is underestimated by LIN FC. (B) Scatterplots of different types of ground truth relationships (linear, nonlinear, or linear and nonlinear) illustrate the theoretical sensitivity of LIN and ENL methods to qualitatively distinct joint distributions at varying levels of added noise. LIN (Pearson correlation) and NULL ENL (NULL) values are sensitive to linearity, but insensitive to nonlinearity. NL (bias-corrected distance correlation) values are sensitive to both linear and nonlinear relationships. ENL values exhibit insensitivity to linearity but sensitivity to nonlinearity. (C) Group-level ICA was implemented in the connectivity domain to obtain spatial representations of networks embedded in the LIN and ENL FC data. Subsequently, the spatial variation of LIN and ENL networks was analyzed. Finally, network associations with corrected postnatal age were assessed using a generalized additive model (GAM) approach. (D) ENL posterior default mode network GAM fit when averaging across 50 voxels with the highest GAM effective degrees of freedom (EDF; quantifies nonlinearity of developmental trajectory). Created in part using BioRender. Kinsey, S. (2026) https://BioRender.com/rrg8drd.

Building on the view that ENL dependence reflects forms of statistical dependence not captured by Gaussian-based (i.e., linear) models (Hlinka et al., 2011), we computed whole-brain FC matrices using distance-correlation, a non-parametric measure sensitive to both linear and nonlinear relationships (Székely & Rizzo, 2014; Székely et al., 2007), and contrasted these with the matrices predicted by a multivariate Gaussian null model (Edelmann et al., 2021). This allowed us to isolate nonlinear FC signatures that violate expectations under assumptions of linear, Gaussian relationships between brain voxels, thereby revealing the ENL patterns that linear methods would overlook (Fig. 1A-B).

To learn network structure directly from this information, we first leveraged group-level spatial independent component analysis (ICA) in the connectivity domain to identify a set of spatially distinct group-level features which serve as proxy representations of the coherent brain ensembles embedded within the data, or intrinsic connectivity networks (ICNs, hereafter referred to as “networks”) (Calhoun et al., 2009; Iraji et al., 2016) (Fig. 1C). We subsequently reconstructed participant-level network maps using the group-level representations as priors. By capitalizing on the explicit separation of nonlinear and linear FC information in addition to the power of connectivity domain ICA to disentangle reliable features from constellations of voxel-wise dependencies (Iraji et al., 2016; Wu et al., 2018), our approach enabled the identification of ENL networks from fMRI data in a highly principled, data-driven manner. Using this method alongside flexible developmental modeling (Feldman et al., 2025), we examined the longitudinal trajectories of ENL and LIN networks (Fig. 1C-D) in a group of infants that were scanned as part of a broader longitudinal study of infants at low and elevated familial genetic likelihood for autism spectrum disorder (ASD) at Marcus Autism Center, Children’s Healthcare of Atlanta. Given ongoing work on neurodevelopmental disorders (Jones & Klin, 2013), the present study focuses on typically developing infants, with “typicality” ascertained on the basis of low familial genetic likelihood for ASD and other neurodevelopmental disability, no history of pre- or perinatal complications, no history of seizures, no known medical conditions or genetic disorders, no hearing loss or visual impairment, and no clinical concerns upon evaluation during toddlerhood (Fig. 2, Table S1).

**Fig. 2.**
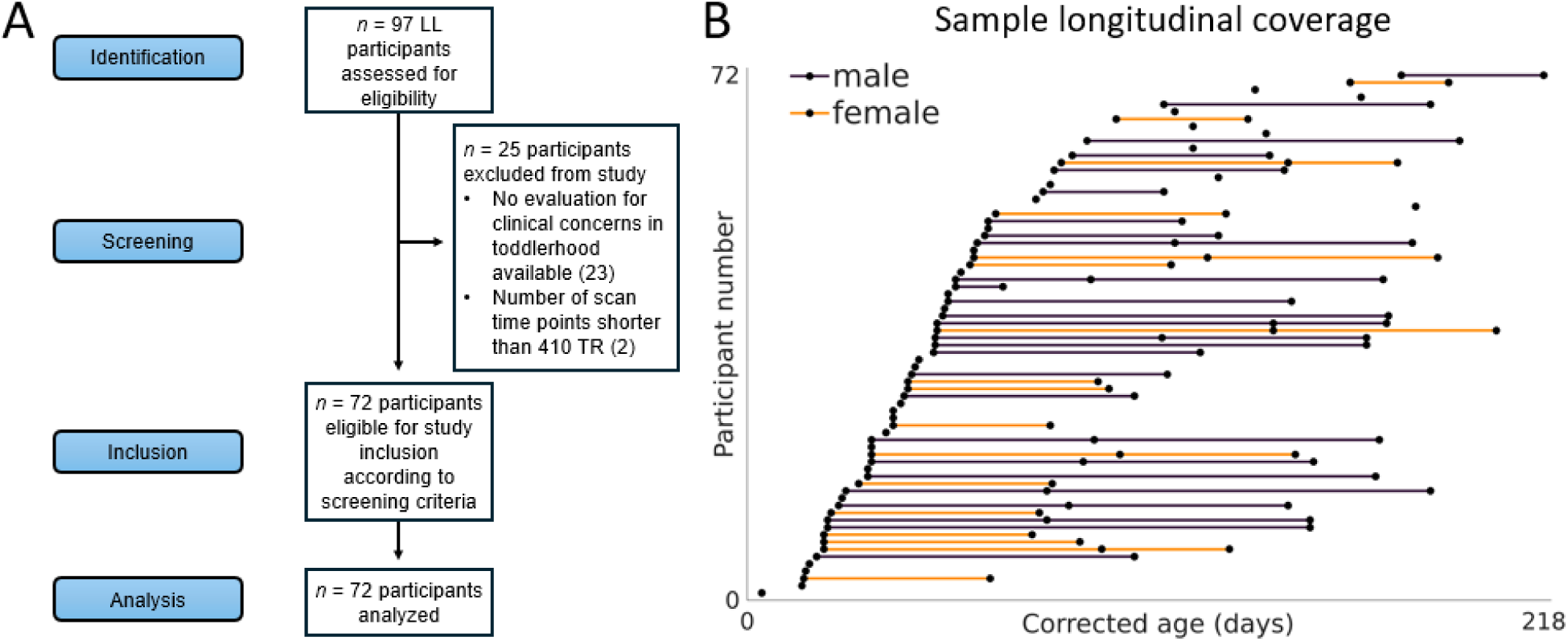
Sample information. (A) Strengthening the reporting of observational studies in epidemiology (STROBE) flowchart. (B) Line plot illustrating the longitudinal coverage of *n* = 130 analyzed resting-state functional magnetic resonance imaging scans. LL: low likelihood for autism spectrum disorder. TR: repetition time.

This framework enabled us to map ENL and LIN network changes during early life, revealing rich longitudinal properties and patterns that have been missed by standard FC methods.

## Results

### Explicitly nonlinear intrinsic connectivity networks are present during early life and reveal differential regional contributions to functional brain connectivity

We extracted 19 ENL networks from the data, indicating that nonlinear fMRI dependencies differentiate brain ensembles as early as birth. Among 16 LIN networks, we identified 14 that were concordant with the ENL approach, suggesting that they strongly contribute to linear and nonlinear FC patterns. We identified no significant difference in the ICASSO quality index (IQ) (Himberg et al., 2004) of concordant ENL and LIN counterparts (*n* = 14 ENL, *n* = 14 LIN; *p* = .9726, difference = −0.000167) or of the full network sets (*n* = 19 ENL, *n* = 16 LIN, *p* = .4037, −0.0064), demonstrating that ENL networks are as reliably estimated as LIN during early postnatal life.

Notably, the data reveal a pronounced primary-to-higher-order gradient in spatial concordance between ENL and LIN (Fig. 3A, graph center). Lower brain structures, primary sensory regions, and motor structures exhibited high concordance, whereas networks comprised of higher-order and associative regions (Margulies et al. 2016, Jensen et al., 2024) exhibited progressively reduced spatial homogeneity. This suggests that higher-order networks are more heterogeneous in terms of their regional contributions to nonlinear and linear FC during infancy. Furthermore, we found that quaternary visual (historically identified as the “where” pathway) (Ungerleider & Mishkin, 1982), left frontoparietal, right frontoparietal, prefrontal, and salience networks did not meet our criteria for concordance and were categorized as unique to the ENL dataset at the given ICA parameters (see Methods), indicating that they have a pronounced nonlinear presence in the infant brain even when they would be missed by linear analysis (Fig. 3B). On the other hand, the left postcentral gyrus and left temporo-parietal-occipital networks were categorized as unique to the LIN dataset, reflecting more prominent contributions to linear dependence patterns (Fig. 3C).

**Fig. 3.**
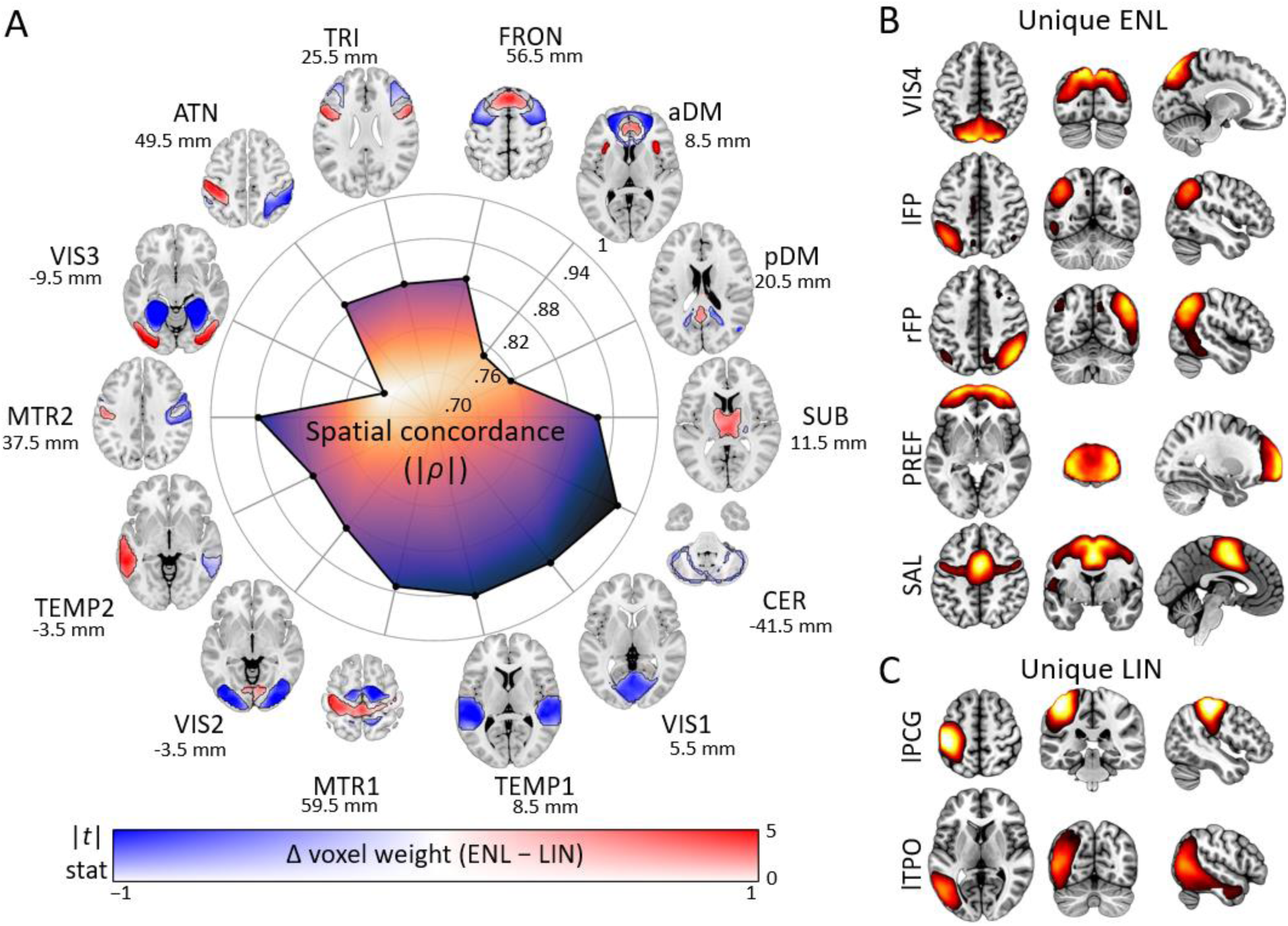
Intrinsic connectivity networks extracted from explicitly nonlinear (ENL) and linear (LIN) functional connectivity. (A) Spatial variation in regional contributions to ENL and LIN FC datasets. Concordant networks (*n* = 14) are displayed around a radial graph where radius indicates degree of spatial concordance (absolute value of correlation) between group-level ENL and LIN counterparts. Concordant networks included subcortical (SUB), cerebellum (CER), primary visual (VIS1), primary temporal (TEMP1), primary sensorimotor (MTR1), secondary visual (VIS2), secondary temporal (TEMP2), secondary sensorimotor (MTR2), tertiary visual (VIS3), dorsal attention (ATN), triangularis (TRI), frontal (FRON), anterior default mode (aDM), and posterior default mode (pDM). For each concordant network pair, maps depict spatial variation between ENL and LIN counterparts according to a dual coded color map, with hue reflecting the difference in voxel weight between ENL and LIN, transparency reflecting *t* statistic magnitude, and contours indicating FDR-corrected significance (*q* < .05). Results are overlaid on the Montreal Neurological Institute (MNI) 152 template with z coordinates listed relative to the origin. (B) Unique ENL networks included quaternary visual (VIS4), left frontoparietal (lFP), right frontoparietal (rFP), prefrontal (PREF), and salience (SAL). (C) Unique LIN networks included left postcentral gyrus (lPCG) and left temporo-parietal-occipital (lTPO). For (B) and (C), results are depicted using an empirical threshold (*Z* > 1.96) overlaid on the MNI 152 template.

We identified significant spatial variation between concordant ENL and LIN counterparts, highlighting differential regional contributions to nonlinear and linear FC data (Fig. 3A, outside graph). A subset of networks including subcortical, secondary visual, triangularis, frontal, anterior default mode, and posterior default mode exhibited core-periphery gradients higher ENL weight in network centers. Another subset including primary sensorimotor, secondary temporal, secondary sensorimotor, and dorsal attention exhibited lateralized patterns. The robustness of our two-sided paired samples *t*-test approach was validated by comparison to the hypothesis outcomes of voxel-wise two-sided permutation tests for the posterior default mode network (ρ = .9930). Summary spatial variation test information can be found in Table S2.

### Explicitly nonlinear and linear intrinsic connectivity networks capture geometrically complex developmental trajectories

Given that ENL networks were just as reliably estimated as LIN, we sought to investigate localized (i.e., voxel-level) associations with corrected age for both methods. Prior work has shown that brain maturation can follow trajectories with varied geometric complexity and that capturing this complexity can reveal important developmental transitions (Faghiri et al., 2019; Feldman et al., 2025; Luna, 2009; Seraji et al., 2025). Therefore, we utilized a flexible generalized additive modeling (GAM) approach and the associated effective degrees of freedom (EDF) to quantify the nonlinearity of developmental trajectories from approximately linear (EDF = 1) to highly nonlinear (EDF ≥ 3) in nature. This allowed us to identify nonlinearities in the contributions of voxels to networks over time, thereby capturing a broad array of developmental associations which would be missed by standard methods. Consequently, our study highlights the informational richness of nonlinear relationships in fMRI data at two distinct levels of analysis.

While we found no significant overall difference in EDF between methods (*p* = .8078, difference = −0.0028), we found that the majority of ENL network counterparts exhibit significantly higher EDF and that the ENL method reveals a larger number of voxels with highly complex (EDF ≥ 3) developmental properties (Table S3). As a case in point, the ENL posterior default mode network develops more nonlinearly than its LIN counterpart (*p* < .001, difference = 0.3515, Hedges’s *g* = 0.3485) (Fig. 4), with high-EDF voxels sharing a longitudinally dynamic pattern of network engagement with rapid increases during the first hundred days of life (Fig. 1D).

**Fig. 4.**
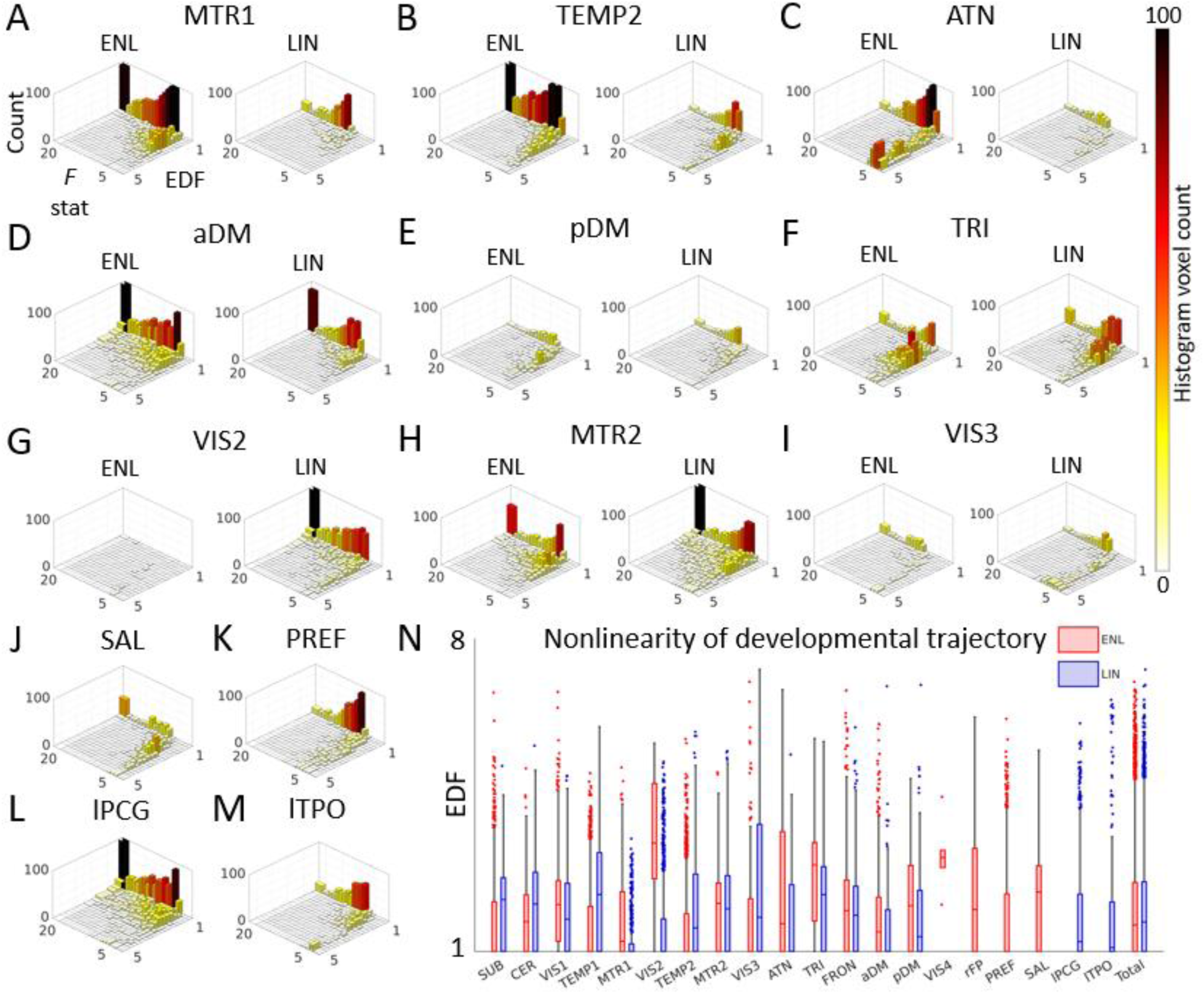
Geometric complexity of explicitly nonlinear (ENL) and linear (LIN) intrinsic connectivity network developmental trajectories. (A-M) Two-dimensional histograms depict age association *F* statistics against effective degrees of freedom (EDF, quantifies nonlinearity of developmental trajectory) for significant ENL and LIN voxels for a representative subset of networks. Hue and bar height represent voxel count. (N) Box and whisker plots depict EDF of significant ENL and LIN voxels. SUB: subcortical. CER: cerebellum. VIS1: primary visual. TEMP1: primary temporal. MTR1: primary sensorimotor. VIS2: secondary visual. TEMP2: secondary temporal. MTR2: secondary sensorimotor. VIS3: tertiary visual. ATN: dorsal attention. TRI: triangularis. FRON: frontal. aDM: anterior default mode. pDM: posterior default mode. VIS4: quaternary visual (ENL). rFP: right frontoparietal (ENL). PREF: prefrontal (ENL). SAL: salience (ENL). lPCG: left postcentral gyrus (LIN). lTPO: left temporo-parietal-occipital (LIN).

We further identified higher EDF for ENL primary visual (*p* < .001, difference = 0.1931, Hedges’s *g* = 0.2195), primary sensorimotor (*p* < .001, difference = 0.4037, Hedges’s *g* = 0.5401), secondary visual (*p* < .001, difference = 2.0974, Hedges’s *g* = 2.5359), dorsal attention (*p* < .001, difference = 0.6296, Hedges’s *g* = 0.4182), triangularis (*p* < .001, difference = 0.5331, Hedges’s *g* = 0.5179), frontal (*p* < .001, difference = 0.1204, Hedges’s *g* = 0.1312), and anterior default mode (*p* < .001, difference = 0.2007, Hedges’s *g* = 0.2666) networks. However, LIN EDF was higher for subcortical (*p* < .001, difference = −0.3831, Hedges’s *g* = 0.4222), cerebellum (*p* < .001, difference = −0.3439, Hedges’s *g* = 0.4175), primary temporal (*p* < .001, difference = −0.7683, Hedges’s *g* = 0.7608), secondary temporal (*p* < .001, difference = –0.3903, Hedges’s *g* = 0.4334), and tertiary visual (*p* < .001, difference = –0.7709, Hedges’s *g* = 0.5117) networks. See Fig. S1 for histograms for remaining networks. See Table S3 for summary age association information and Table S4 for summary EDF comparison information.

### Explicitly nonlinear intrinsic connectivity networks reveal hidden aspects of infant brain development

Upon localizing developmental associations, we found that a substantial subset of ENL networks reflect spatially expansive developmental patterns (Fig. 5). Among concordant networks, the ENL counterparts of primary sensorimotor (Fig. 5A), secondary temporal (Fig. 5B), dorsal attention (Fig. 5C), anterior default mode (Fig. 5D), posterior default mode (Fig. 5E), and triangularis (Fig. 5F) revealed voluminous clusters missed by the LIN method. For example, the ENL primary sensorimotor network exhibits nonlinear developmental trajectories across the postcentral gyri and paracentral lobules, with linear trajectories within the precuneus. On the other hand, LIN primary sensorimotor trajectories were mostly linear and confined to the right precentral gyrus. Furthermore, we found that ENL anterior and posterior default mode networks develop nonlinearly within core regions such as the anterior and posterior cingulum, respectively. In contrast, LIN default mode development is more sparse and localized mainly to peripheral regions. The ENL counterpart of the triangularis network revealed hidden, geometrically complex (EDF ≥ 3) developmental patterns in the bilateral insula and frontal inferior operculum. On the other hand, we note that the LIN method detected sizable clusters in secondary (Fig. 5G) and tertiary (Fig. 5I) visual networks that were missed by the ENL method.

**Fig. 5.**
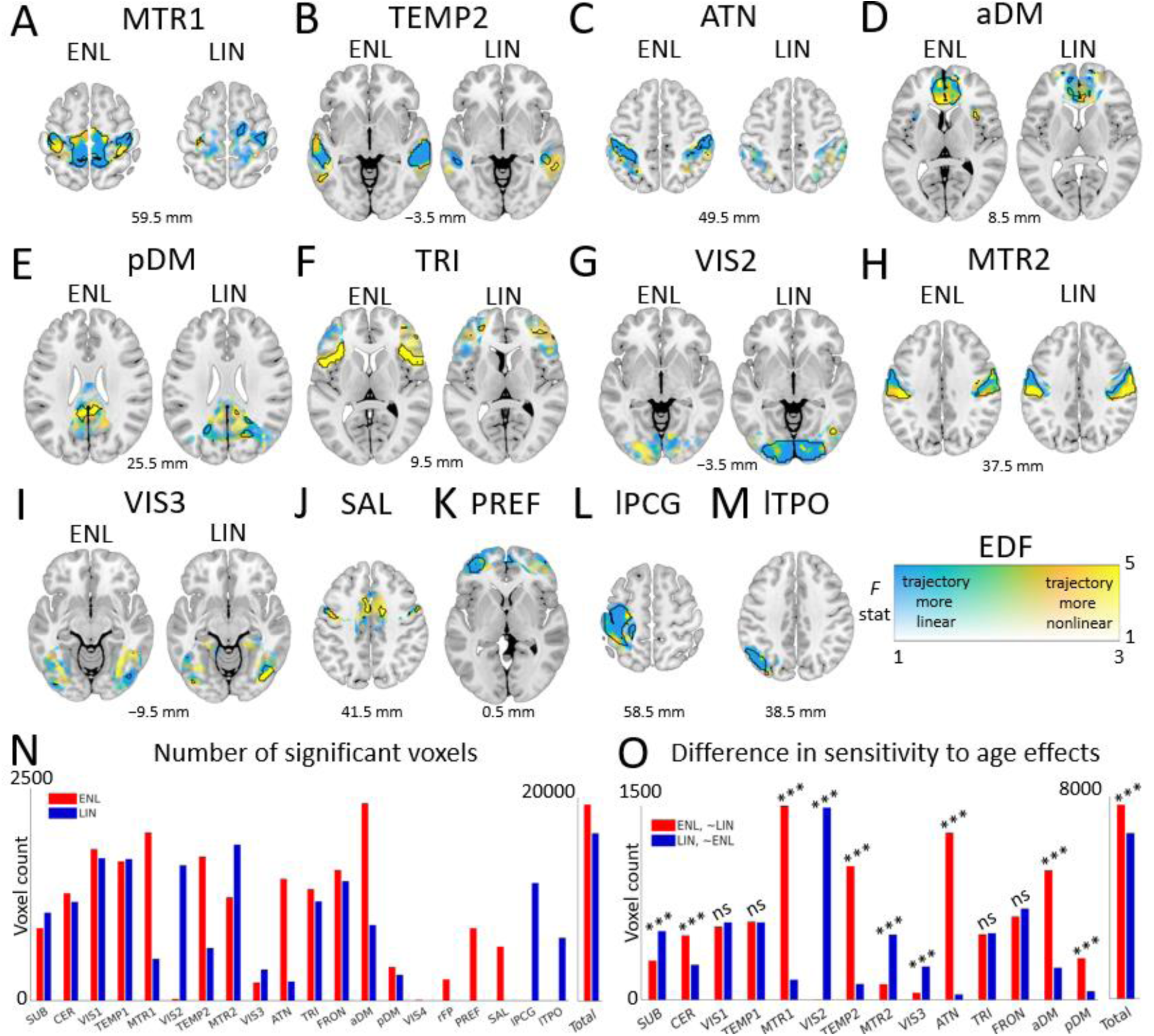
Localization and sensitivity of explicitly nonlinear (ENL) and linear (LIN) intrinsic connectivity network associations with corrected postnatal age. (A-M) Spatial maps of voxel associations with age are plotted according to a dual-coded colormap for a representative subset of ENL and LIN networks, with transparency reflecting *F* statistic magnitude, hue representing model effective degrees of freedom (EDF; quantifies nonlinearity of developmental trajectory), and black contours indicating FDR-corrected significance (*q* < .05). Warmer hues indicate more complex, nonlinear developmental trajectories while cooler hues indicate less complex, approximately linear trajectories. Results are overlaid on the Montreal Neurological Institute 152 template with z coordinates listed relative to the origin. (N) Number of significant voxels identified using ENL and LIN methods. (O) Differences in the statistical sensitivity of ENL and LIN methods to age effects highlight the relevance of nonlinear fMRI relationships to brain development. SUB: subcortical. CER: cerebellum. VIS1: primary visual. TEMP1: primary temporal. MTR1: primary sensorimotor. VIS2: secondary visual. TEMP2: secondary temporal. MTR2: secondary sensorimotor. VIS3: tertiary visual. ATN: dorsal attention. TRI: triangularis. FRON: frontal. aDM: anterior default mode. pDM: posterior default mode. VIS4: quaternary visual (ENL). rFP: right frontoparietal (ENL). PREF: prefrontal (ENL). SAL: salience (ENL). lPCG: left postcentral gyrus (LIN). lTPO: left temporo-parietal-occipital (LIN).

With respect to uniquely identified networks, we found that ENL salience (Fig. 5J) and prefrontal (Fig. 5K) revealed prominent developmental patterns. However, we also detected large clusters in the LIN left postcentral gyrus (Fig. 5L) and left temporo-parietal-occipital (Fig. 5M) networks. See Fig. S2 for developmental maps of remaining networks. See Table S3 for summary age association information.

### Explicitly nonlinear intrinsic connectivity networks exhibit enhanced sensitivity to age effects

Overall, the ENL counterparts of the concordant networks showed markedly greater statistical sensitivity to corrected age relative to LIN counterparts (𝜒^2^ = 353.4705, *p* < .00001, odds ratio (OR) = 1.39) (Fig. 5O, Table S5), indicating that features derived from nonlinear FC may serve as more informative neurodevelopmental biomarkers. Furthermore, individual comparisons of concordant network pairs revealed a statistical advantage for ENL primary sensorimotor (𝜒^2^ = 1019.5, *p* < .00001, OR = 9.77), dorsal attention (𝜒^2^ = 1090.6, *p* < .00001, OR = 32.41), anterior default mode (𝜒^2^ = 421.9352, *p* < .00001, OR = 4.06), posterior default mode (𝜒^2^ = 156.0112, *p* < .00001, OR = 4.95), cerebellum (𝜒^2^ = 62.0253, *p* < .00001, OR = 1.84), and secondary temporal (𝜒^2^ = 668.2689, *p* < .00001, OR = 8.56) networks. While an advantage for the LIN method was discovered for subcortical (𝜒^2^ = 57.6697, *p* < .00001, OR = 1.76), secondary visual (𝜒^2^ = 1375, *p* < .00001, OR = 1380), tertiary visual (𝜒^2^ = 123.1498, *p* < .00001, OR = 4.86), and secondary sensorimotor (𝜒^2^ = 218.0363, *p* < .00001, OR = 4.21) networks, this collection of findings strongly underwrites the relevance of nonlinear fMRI connectivity to studies of brain development. See Table S5 for summary statistical sensitivity test information.

## Discussion

For the first time, our study establishes that fMRI ensembles systematically participate in nonlinear relationships as early as birth. Surprisingly, our findings further indicate that perhaps the bulk of infant brain network development goes undetected by linear approaches. In contrast with studies which suggest that nonlinear fMRI FC is unreliable (Raffaelli et al., 2024), or fails to provide an incremental advantage over linear methods in detecting age associations (Jin et al., 2025), our findings collectively underscore its informational richness and indicate that the insights it affords can go overlooked because of methodological decisions made by researchers. Our study supports the view that applications of nonlinear estimation methods at the macroscale should be assessed in terms of their potential to capture aspects of the brain’s neurofunctional organization, as even in cases where nonlinear fMRI FC patterns account for a fraction of the total dependence, they may reflect important biophysical properties of the neural populations underlying the BOLD signal. This underscores the importance of expanding beyond assumptions of linearity to capture the full structure of functional brain interactions.

With respect to patterns of spatial variation between network counterparts, our results are largely in line with our previous finding that large-scale brain networks exhibit a core-periphery gradient in terms of their contributions to ENL and LIN FC information (Kinsey et al., 2024). However, networks including primary sensorimotor, secondary temporal, secondary sensorimotor, and dorsal attention exhibited more lateralized regional contributions to nonlinear and linear FC in the infant dataset analyzed here. These findings suggest that, for some brain ensembles, lateralized patterns of nonlinear and linear FC contributions shift toward a bilateral center-periphery organization over the course of maturation. Moreover, the detection of unique networks such as salience, left frontoparietal, right frontoparietal, and prefrontal from ENL (but not LIN) data (Fig. 3B) suggests that they have a pronounced nonlinear presence during early life even when they would be missed by standard estimation methods. Such findings are supported by prior research linking functional brain nonlinearity to limbic regions in humans (Kinsey et al., 2024; Jin et al., 2025) and to prefrontal regions in primates (Rigotti et al., 2013; Johnston et al., 2020; Fusi et al., 2016), although our findings implicate prefrontal regions in nonlinear relationships earlier in the developmental cycle and at several greater orders of spatiotemporal magnitude.

Our developmental analysis motivates the investigation of the role of nonlinear relationships in the development of a wide range of cognitive capacities. Broadly speaking, the relatively greater degree of developmental nonlinearity for multiple ENL counterparts indicates that nonlinear FC is a locus of valuable information about temporally localized maturation epochs in the infant brain. While the development of ENL primary sensorimotor connectivity may capture aspects of the refinement of motor capacities during the earliest stages of life, the development of ENL posterior default mode connectivity may likewise capture proto-development in cognitive capacities such as self-referential ideation and social cognition (Wang et al., 2020). Furthermore, the ENL counterpart of the triangularis network exhibited geometrically complex trajectories within insular regions that have been previously associated with linguistic capacities (Oh et al., 2014). Considering that this network also integrated regions such as Broca’s area (Fridrikkson et al., 2014), our findings suggest that changes in this network’s nonlinear FC patterns may contribute to foundational, temporally localized developments in the capacity for language expression. Additionally, the salience network revealed similarly complex developmental profiles within middle cingulate regions associated with stimulus salience and social reward (Apps et al., 2013) in addition to trajectories within precentral gyrus regions that are key to motor function (Fig. 5J), indicating the possibility that the development of social reward processing is underwritten by the longitudinal functional differentiation of these regions. With respect to more domain-general cognitive capacities, the left lateralized developmental patterns of ENL weight identified in the prefrontal network (Fig. 5K) may reflect refinements in the flexible computational properties (Fusi et al., 2016) of these regions.

From a methodological perspective, some studies have investigated non-Gaussian dependence in fMRI data using mutual information–based approaches (Hlinka et al., 2011; Raffaelli et al., 2024). Mutual information is sensitive to nonlinear associations and is often regarded as relatively equitable, meaning that it tends to assign similar dependence values to relationships that have comparable noise levels, regardless of whether the underlying form is linear, quadratic, periodic, or otherwise (Kinney et al., 2014). Here, we estimate ENL FC using a distance correlation framework. Unlike mutual information, distance correlation is inherently non-equitable, such that it often yields different dependence magnitudes for relationships that share the same noise level but differ in their functional form. This selective sensitivity means that distance-correlation–based ENL estimates naturally differentiate among distinct types of nonlinear interactions. As a result, our approach offers potentially enhanced ability to distinguish between functional brain ensembles that participate in qualitatively distinct nonlinear relationships and perform fundamentally different types of internal signal transformations (Rathkopf, 2013).

Although our study marks an important step forward in neurodevelopmental biomarker construction, it has several limitations. First, due to the limited sample size of the present study, larger longitudinal studies are required to further validate the identified developmental patterns. However, this consideration does not bear on our comparison between the statistical sensitivity of network estimation methods at the given sample size. Second, our conclusions are limited to populations with demographic characteristics similar to the analyzed sample and cannot necessarily be generalized to populations that exhibit distinct demographic characteristics. Third, because our study utilized an ICA model order designed to capture large-scale networks (see Methods), we leave investigations of more localized, fine-grained networks (Iraji et al., 2022; Iraji et al., 2023b; Mirzaeian et al., 2025) to future studies. Fourth, while we identified networks that were unique to LIN and ENL methods (Fig. 3B-C), we cannot rule out the possibility that they might have been identified from both datasets with adjustments of ICA parameters, such as model order (Iraji et al., 2023b; Bajracharya et al., 2024). Nevertheless, our findings emphasize the relative presence of these networks within their respective datasets and highlight the information that is lost by standard analyses when parameters are held constant. Fifth and finally, while we emphasize the rich information carried by nonlinear fMRI connectivity in our study, our aim is not to relegate linear connectivity patterns. Indeed, our findings indicate that linear and nonlinear methods both yield valuable information about functional brain connectivity. In summary, our work serves as a landmark for future studies of brain complexity and development by emphasizing the richness of typically neglected nonlinear relationships in fMRI data and their prospective utility in early screening for neurodevelopmental and psychiatric risk.

## Methods

### Participant information and data acquisition

We analyzed resting-state fMRI scans acquired from typically developing infants as part of a study conducted by the Marcus Autism Center, i/o Children’s Healthcare of Atlanta. Parents provided informed written consent and all regulations relevant to human research participants were followed. The Emory University Institutional Review Board approved the research protocol. Participants were screened for developmental delays in first-degree relatives, history of ASD in up to third-degree relatives, pre- and perinatal complications, history of seizures, medical conditions, genetic disorders, hearing or visual impairments, contraindications for MRI, and clinical concerns when evaluated in toddlerhood. Each infant in the broader study was scanned at up to three pseudorandom time points between birth and 287 postnatal days. All data were collected using either a Siemens TIM Trio scanner (repetition time (TR) = 720 ms; echo time (TE) = 33 ms) or a Siemens MAGNETOM Prisma scanner (TR = 800 ms; TE = 37 ms). We screened scans collected from potentially eligible infants for data length suitable for group-level ICA, as low scan length and/or high length heterogeneity between participants and sessions have been identified as confounds for such analyses (Calhoun et al., 2008; Duda et al., 2023; Iraji et al. 2023b). The vast majority of potentially eligible scans met a length threshold of 410 TR. Therefore, we used this TR threshold when selecting scans for further analysis. Because scans were phase encoded in AP or PA directions, we controlled the impact of phase encoding direction on subsequent analyses by selecting one AP scan per session at random, resulting in a final pool of *n* = 130 scans collected from *n* = 72 infants. The corrected age for each scan was calculated to adjust for preterm or post term effects and was used as the predictor of interest in developmental models (Seraji et al., 2025). We controlled for sex as a biological variable via its inclusion as a term in developmental models.

### Preprocessing

Initial preprocessing steps included the removal of the first ten volumes to ensure magnetization equilibrium, susceptibility distortion correction using FSL (https://fsl.fmrib.ox.ac.uk/fsl/fslwiki) with high-resolution reference images, slice timing correction, rigid body motion realignment implemented in SPM (https://www.fil.ion.ucl.ac.uk/spm/), two-stage normalization to Montreal Neurological Institute (MNI) 152 coordinate space consisting of initial registration to a four-month infant template derived from the Baby Connectome Project cohort (Chen et al., 2022) followed by transformation to the MNI 152 echo-planar imaging template, and spatial smoothing with a 6 mm full width at half maximum (FWHM) Gaussian kernel. Additional preprocessing and cleaning steps were performed within the MATLAB (R2020b) software environment including head motion regression, detrending, despiking, low pass filtering, temporal resampling to TR = 720 ms, and voxel time series *Z*-scoring to normalize variance.

### Estimating explicitly nonlinear and linear fMRI functional connectivity

We previously estimated ENL FC by removing Pearson correlation information from distance correlation via whole-brain FC regression (Kinsey et al., 2024). However, we did not directly account for the distinctness of the Pearson correlation and distance correlation metric spaces (Székely et al, 2007; Edelmann et al., 2021). Furthermore, our earlier approach did not directly account for statistical confounds including sample length bias (Monroy-Castillo et al., 2025; Székely & Rizzo, 2014). Here, we address these limitations by computing metric space-transformed, bias-corrected ENL FC matrices. This framework allows principled and precise access to observed distance correlation patterns that supersede expectations under the assumption of fMRI multivariate Gaussianity.

Let 𝑋 ∈ ℝ^𝑛×𝑣^be a sample of fMRI data where 𝑛 is the number of time points, 𝑣 is the number of voxels within the brain, and 𝑥 and 𝑦 represent any two preprocessed voxel time series such that 𝑥, 𝑦 ∈ ℝ^1×𝑛^. Thus, 𝑥_𝑖_ is the value of voxel 𝑥 at time point 𝑖. First, we calculated the squared bias-corrected distance correlation (Monroy-Castillo et al., 2025; Székely & Rizzo, 2014) (ℛ^2^) (Eq. 1) across all pairs of brain voxels to capture nonlinear statistical dependencies. The squared bias-corrected distance correlation is defined as

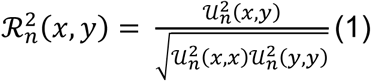

where the squared unbiased distance covariance (𝒰^2^) is

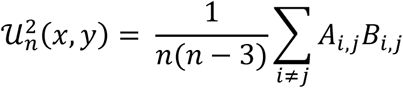

and 𝐴 and 𝐵 represent 𝒰-centered versions of Euclidean distance matrices for voxel time series 𝑥 and 𝑦 such that

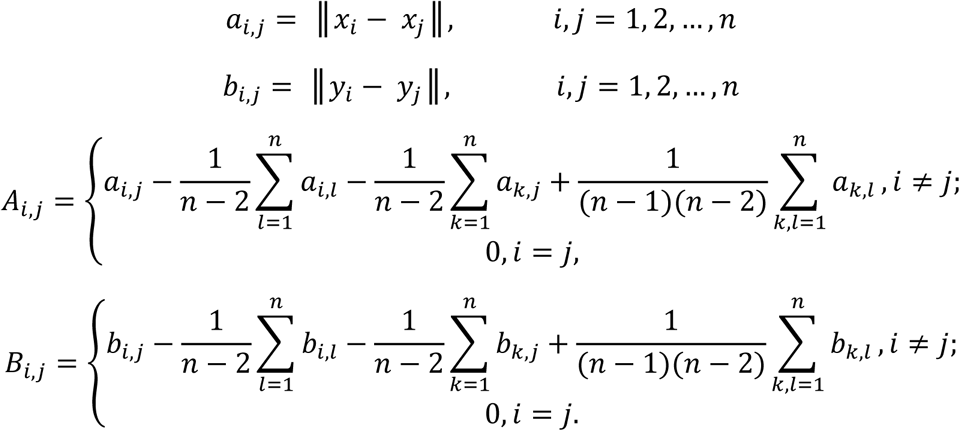

The original distance correlation estimator is known to be positively biased for smaller sample sizes, resulting in the overestimation of statistical dependence between variables that exhibit ground truth dependence equal to or near zero (Monroy-Castillo et al., 2025; Székely & Rizzo, 2014). Therefore, we utilized the bias-corrected distance correlation estimator to directly mitigate any dependence bias resulting from sample length (Monroy-Castillo et al., 2025).

Because the squared bias-corrected distance correlation can take negative values (Monroy-Castillo et al., 2025; Székely & Rizzo, 2014), several solutions have been proposed to deal with the problem of estimating bias-corrected distance correlation. For this study, we truncated all negative values to zero and took the square root of the truncated values, as this computationally straightforward approach has been shown to perform well under a variety of simulated conditions (Monroy-Castillo et al., 2025). Thus, we estimated the nonlinear (NL) FC for each scan per Eq. 2.

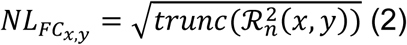

We note that distance correlation is sensitive to both linear and nonlinear relationships. To remove the linear FC information, we first calculated the Pearson correlation coefficient (𝑟) across voxel time series pairs (Eq. 3).

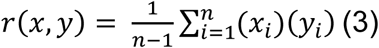

We then transformed these values into the distance correlation space under the assumption of bivariate Gaussianity (Edelmann et al., 2021), yielding a NULL ENL FC matrix for each rsfMRI scan (Eq. 4).

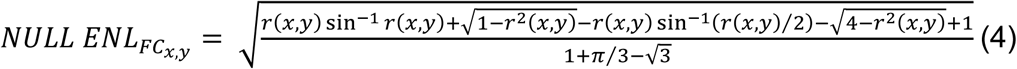

Finally, we subtracted NULL ENL FC from NL FC, yielding whole-brain ENL FC (Eq. 5).

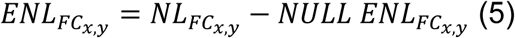

NULL ENL FC represents the counterfactual distance correlation – what the distance correlation would be if voxel time series 𝑥 and 𝑦 were jointly distributed as bivariate Gaussian and nonlinear effects were absent or negligible. Therefore, the ENL FC quantifies the statistical dependence between 𝑥 and 𝑦 which supersedes the linear dependence (i.e., the ENL dependence). Importantly, although prior studies have cited the computational scalability of nonlinear methods as a drawback (Jin et al., 2025; Raffaelli et al., 2024), we provide ENL FC code in vectorized form, which allows for efficient whole-brain analyses of large numbers of participants. Our method is therefore highly scalable. See code availability for more details.

### Extracting intrinsic connectivity networks

The group ICA of fMRI Toolbox (GIFT v4.0; http://trendscenter.org/software/gift) (Iraji et al., 2021) was used to estimate separate sets of networks from the LIN FC and ENL FC data. Principal component analysis (PCA) was first implemented to denoise and reduce the dimensionality of the FC matrices estimated from each scan. The 30 principal components (PCs) that explained the maximum variance of each matrix were retained. All PCs were then concatenated, an additional group-level PCA step was performed retaining the top 20 group-level PCs, and blind group-level spatial independent component analysis (gr-sICA) was implemented in the connectivity domain via the Infomax (Bell & Sejnowski, 1995) optimization algorithm 100 times with random initialization and bootstrapping to increase the chances of escaping local minima. We selected a model order of 20 to obtain large-scale networks, and we used the ICASSO IQ to assess component estimation reliability (Himberg et al., 2004). A component was classified as a network if and only if 1) it exhibited peak weight in and visual overlap with gray matter, 2) it exhibited low visual similarity to known artifacts, and 3) it exhibited an ICASSO IQ value exceeding .80. Components were matched in a greedy fashion based on the absolute value of their spatial correlation, and matched networks were visually inspected and received concordant categorical labels based on their neuroanatomical distributions and recommendations from prior studies (Iraji et al., 2016; Iraji et al., 2023b). Specifically, a network pair was categorized as concordant if ENL and LIN counterparts exhibited absolute value of spatial correlation greater than .70 and were labeled identically based on prior expertise. To assess differences in estimation reliability between ENL and LIN concordant network counterparts, we conducted two-sided paired samples permutation tests with 5000 random permutations. To assess differences in estimation reliability between the full ENL and LIN network sets separately, we conducted two-sided independent samples permutation tests with 5000 random permutations (Krol, 2023; https://github.com/lrkrol/permutationTest). We then implemented group information-guided ICA (Du & Fan, 2013) via GIFT to reconstruct participant-level spatial maps from the group-level networks and participant-level PCs.

### Assessing spatial variation between concordant explicitly nonlinear and linear intrinsic connectivity network counterparts

To assess spatial variation between concordant networks, we conducted two-sided paired samples *t*-tests on the participant-level weights of voxels that significantly contributed to either group-level map (*Z* > 1.96 ENL ∨ LIN empirical threshold). The Benjamini-Hochberg (Benjamini & Hochberg, 1995) false discovery rate (FDR) method was used to correct for multiple comparisons across voxels. The robustness of the *t*-test procedure was assessed via comparison to the FDR-corrected hypothesis outcomes of two-sided paired samples permutation tests with 5000 random permutations for the posterior default mode network.

### Identifying voxel-level associations with corrected age

To assess relationships between voxel contributions to networks and postnatal age, we utilized a GAM approach which flexibly accommodates both linear and nonlinear developmental trajectories (Hastie & Tibshirani, 1990). In the context of developmental analyses, prior work has demonstrated the value of analyzing the nonlinear trajectory of independent variables such as behavioral factors (Luna, 2009) and network properties (Faghiri et al., 2019; Feldman et al., 2025; Seraji et al., 2025), necessitating methods that move beyond a traditional general linear model. GAM extends the generalized linear model by allowing smooth, potentially nonlinear functions of predictors to be learned directly from the data, providing a principled framework for modeling age-related effects without imposing restrictive parametric assumptions. For each LIN and ENL network, we used a GAM approach to identify associations between the participant-level weights of voxels that significantly contributed to their respective group-level maps (*Z* > 1.96 empirical threshold) (dependent variable) and corrected age (independent variable) while controlling for confounders including participant ID (random effect), sex, scanner, and motion (mean framewise displacement) (fixed effects). For each network, Benjamini-Hochberg FDR correction was used to correct for multiple comparisons across brain voxels. The automated anatomical labeling atlas 3 (AAL3) (Rolls et al., 2020) was used when localizing significant voxels to anatomically defined brain regions.

To assess differences in developmental nonlinearity for ENL and LIN methods, we conducted two-sided independent samples permutation tests with 5000 random permutations on the EDF values of significant voxels for concordant networks individually in addition to a separate test on the EDF values of all significant voxels. The Bonferroni family-wise error method was used to correct for multiple comparisons (*p* < .05/15).

### Assessing differences between explicitly nonlinear and linear intrinsic connectivity network sensitivity to corrected age

For concordant network pairs, we first identified the voxels that were assessed for associations with age using both methods. Results for each voxel could fall into one of four outcomes: significant for both methods, significant for LIN only, significant for ENL only, or significant for neither method. We conducted two-sided McNemar’s tests on this outcome information to compare ENL and LIN statistical sensitivity to age for individual network pairs. Finally, we conducted a two-sided McNemar’s test between ENL and LIN methods for all voxels that were collectively tested across concordant network pairs to compare overall statistical sensitivity to age. The Bonferroni family-wise error method was used to correct for multiple comparisons (*p* < .05/15).

## Data availability

Data collected from National Institute of Mental Health (NIMH) 2P50MH100029, R01MH118285, and R01MH119251 are available from the NIMH Data Archive (NDA). NDA is a collaborative informatics system created by the National Institutes of Health to provide a national resource to support and accelerate research in mental health. Data will be released in a NDA study-specific repository upon peer-reviewed publication. The manuscript reflects the views of the authors and may not reflect the opinions or views of the NIH.

## Code availability

Preprocessing and data analysis were conducted primarily within the MATLAB software environment mainly using MATLAB 9.9.0.1857802 (R2020b) Update 7, the Statistical Parametric Mapping toolbox (SPM 12), the FMRIB software library (FSL v6.0), the Group ICA of fMRI toolbox (GIFT v4.0) and RStudio 2025.05.0+496 version (R 4.4.3). MATLAB R2020b can be downloaded from https://www.mathworks.com. The FSL v6.0 toolbox can be downloaded from https://fsl.fmrib.ox.ac.uk/fsl/fslwiki. The SPM 12 toolbox can be downloaded from https://www.fil.ion.ucl.ac.uk/spm/. GIFT v4.0 can be downloaded from https://trendscenter.org/software/gift/. R v4.4.3 can be downloaded from https://cran.r-project.org/. The sample scripts utilized for dual code data visualization can be downloaded from https://trendscenter.org/x/datavis/. The independent samples permutation test function utilized for ICASSO IQ and EDF statistical randomization analyses can be downloaded from https://github.com/lrkrol/permutationTest/ (Krol, 2023). The MATLAB function used to calculate ENL FC will be deposited on a GitHub repository prior to peer-reviewed publication. Other MATLAB code used for this study can be obtained from the corresponding authors upon reasonable request.

## Author contributions

S.K., V.D.C., and A.I. proposed the study. S.S. acquired the original data. Z.F. preprocessed the data. M.S. assisted with data screening. S.K., V.D.C., and A.I. contributed to methods development. S.K., G.N., and P.B. analyzed the data. S.K., A.I., and V.D.C. contributed to the interpretation of the results. S.K. drafted the paper. M.S., V.D.C., S.S., and A.I. contributed to the critical revision of the paper.

## Acknowledgements.

We greatly appreciate the families and their infants who volunteered to participate in this research study. We would also like to thank the research coordinators, assistants, and fellows at the Marcus Autism Center, Brittney Sholar, Carly Reineri, Joannna Beugnon, Lindsey Evans, Jordan Pincus, Jennifer Gutierrez, Tristan Ponzo, and Adriana Mendez, and the MRI techs at the Emory Center for Systems Imaging Core, Michael White, Sarah Basadre, and Samira Yeboah, for their data collection efforts.

The National Institute of Mental Health, U.S.A. (K01MH108741, 2P50MH100029, R01MH118285 to S.S., R01MH119251 to S.S. and A.I., R01MH136665 to A.I.); the National Institute of Biomedical Imaging and Bioengineering, U.S.A. (R01EB027147 to S.S. and V.D.C.); the National Science Foundation, U.S.A. (2112455 to V.D.C.), funds from the Whitehead and Marcus Foundations (to S.S., A.I., and V.D.C.).

