## Supplementary Information for "Explicitly nonlinear fMRI networks reveal hidden trajectories of infant brain development"

### Supplementary Information for Kinsey et al. “Explicitly nonlinear networks reveal hidden trajectories of infant brain development”

**Table S1.** Sample demographic information. For demographic categories with a participant number less than the total sample, *n* is specified in the category title. SD: standard deviation. rsfMRI: resting-state functional magnetic resonance imaging. GED: general educational development.

| **Sex (no.)** | **Race (no.), *n* = 64** | **Maternal education (no.), *n* = 62** | **Household income in United States dollars (no.), *n* = 61** | **Gestational age at birth in weeks (mean ± SD)** | **No. rsfMRI scans** |
|---|---|---|---|---|---|
| Male (42) | White (33)<br>Black or African American (3)<br>More than one race (1) | High school or GED (1)<br>Trade/vocational training (0)<br>College coursework (1)<br>Associate’s degree (1)<br>College degree (13)<br>Master’s degree (16)<br>Professional degree (2) | < $40000 (1)<br>$40000-$60000 (5)<br>$60001-$80000 (2)<br>$80001-$100000 (7)<br>$100001-$125000 (6)<br>$125001-$150000 (4)<br>> $150000 (9) | 39.17 ± 1.46 | 78 |
| Female (30) | White (23)<br>Black or African American (2)<br>More than one race (2) | High school or GED (0)<br>Trade/vocational training (1)<br>College coursework (3)<br>Associate’s degree (0)<br>College degree (9)<br>Master’s degree (11)<br>Professional degree (4) | < $40000 (2)<br>$40000-$60000 (2)<br>$60001-$80000 (3)<br>$80001-$100000 (7)<br>$100001-$125000 (4)<br>$125001-$150000 (4)<br>> $150000 (5) | 39.18 ± 1.16 | 52 |

9

**Table S2.** Summary information for assessment of spatial variation between concordant explicitly nonlinear (ENL) and linear (LIN) intrinsic connectivity network (ICN) counterparts. Statistical tests were voxel-wise two-sided paired samples *t*-tests. Confidence interval (CI) information is reported as mean ± std CI range, while summary effect size information is reported as Cohen's *D* mean ± standard deviation (SD). rsfMRI: resting-state functional magnetic resonance imaging. SUB: subcortical. CER: cerebellum. VIS1: primary visual. TEMP1: primary temporal. MTR1: primary sensorimotor. VIS2: secondary visual. TEMP2: secondary temporal. MTR2: secondary sensorimotor. VIS3: tertiary visual. ATN: dorsal attention. TRI: triangularis. FRON: frontal. aDM: anterior default mode. pDM: posterior default mode.

| **ICN** | ***n* (rsfMRI scans)/*df*** | **CI range (mean ± std) - ENL > LIN** | **CI range (mean ± SD) - LIN > ENL** | **Cohen's *D* (mean ± SD) ENL > LIN** | **Cohen's *D* (mean ± SD) LIN > ENL** |
|---|---|---|---|---|---|
| SUB | 130/129 | 0.3250 ± 0.0303 | 0.3602 ± 0.0311 | 0.4790 ± 0.1934 | 0.5533 ± 0.2141 |
| CER | 130/129 | 0.4211 ± 0.0365 | 0.3894 ± 0.0457 | 0.2201 ± 0.0175 | 0.3516 ± 0.1145 |
| VIS1 | 130/129 | 0.3481 ± 0.0286 | 0.3781 ± 0.0340 | 0.2430 ± 0.0440 | 0.5083 ± 0.1886 |
| TEMP1 | 130/129 | 0.3671 ± 0.0306 | 0.3990 ± 0.0415 | 0.2819 ± 0.0689 | 0.6017 ± 0.2361 |
| MTR1 | 130/129 | 0.4049 ± 0.0356 | 0.3534 ± 0.0442 | 0.4188 ± 0.1749 | 0.5708 ± 0.2931 |
| VIS2 | 130/129 | 0.4303 ± 0.0343 | 0.4031 ± 0.0477 | 0.4911 ± 0.1846 | 0.6326 ± 0.2405 |
| TEMP2 | 130/129 | 0.3479 ± 0.0358 | 0.3331 ± 0.0322 | 0.6265 ± 0.2035 | 0.5895 ± 0.2960 |
| MTR2 | 130/129 | 0.3690 ± 0.0371 | 0.3671 ± 0.0425 | 0.4254 ± 0.1709 | 0.5016 ± 0.2081 |
| VIS3 | 130/129 | 0.4122 ± 0.0357 | 0.3695 ± 0.0320 | 0.8121 ± 0.3376 | 1.1214 ± 0.5010 |
| ATN | 130/129 | 0.3937 ± 0.0351 | 0.3522 ± 0.0431 | 0.4601 ± 0.1595 | 0.6030 ± 0.3036 |

| | | | | | |
|---|---|---|---|---|---|
| TRI | 130/129 | 0.3260 ± 0.0290 | 0.3229 ± 0.0273 | 0.5333 ± 0.2178 | 0.6506 ± 0.3045 |
| FRON | 130/129 | 0.3514 ± 0.0291 | 0.3378 ± 0.0400 | 0.5861 ± 0.2443 | 0.5601 ± 0.2512 |
| aDM | 130/129 | 0.3515 ± 0.0341 | 0.3419 ± 0.0303 | 0.7082 ± 0.3729 | 0.6595 ± 0.2705 |
| pDM | 130/129 | 0.3410 ± 0.0288 | 0.3248 ± 0.0304 | 0.3575 ± 0.1241 | 0.7156 ± 0.3250 |

**Table S3.** Summary information for intrinsic connectivity network (ICN) associations with age. Statistical tests were generalized additive model *F* tests. Summary model effective degrees of freedom (EDF; quantifies nonlinearity of developmental trajectory) information for all tested voxels is reported as mean ± standard deviation (SD). ENL: explicitly nonlinear. LIN: linear. rsfMRI: resting-state functional magnetic resonance imaging. SUB: subcortical. CER: cerebellum. VIS1: primary visual. TEMP1: primary temporal. MTR1: primary sensorimotor. VIS2: secondary visual. TEMP2: secondary temporal. MTR2: secondary sensorimotor. VIS3: tertiary visual. ATN: dorsal attention. TRI: triangularis. FRON: frontal. aDM: anterior default mode. pDM: posterior default mode. VIS4: quaternary visual. lFP: left frontoparietal. rFP: right frontoparietal. PREF: prefrontal. SAL: salience. lPCG: left postcentral gyrus. lTPO: left temporo-parietal-occipital.

| **ICN** | **No. significant voxels - ENL** | **No. significant voxels - LIN** | ***n* (rsfMRI scans)** | **EDF (mean ± SD) - ENL** | **EDF (mean ± SD) - LIN** | **No. voxels EDF ≥ 3 - ENL** | **No. voxels EDF ≥ 3 - LIN** |
|---|---|---|---|---|---|---|---|
| SUB | 835 | 1014 | 130 | 1.6557 ± 1.0177 | 2.0388 ± 0.8054 | 99 | 95 |
| CER | 1239 | 1139 | 130 | 1.7182 ± 0.7207 | 2.0621 ± 0.9227 | 50 | 188 |
| VIS1 | 1748 | 1644 | 130 | 2.0497 ± 0.8631 | 1.8566 ± 0.8963 | 199 | 206 |
| TEMP1 | 1606 | 1634 | 130 | 1.5728 ± 0.8672 | 2.3411 ± 1.1326 | 157 | 483 |
| MTR1 | 1941 | 480 | 130 | 1.6753 ± 0.7890 | 1.2716 ± 0.5487 | 125 | 8 |
| VIS2 | 16 | 1562 | 130 | 1.5199 ± 0.8462 | 1.4767 ± 0.8190 | 11 | 101 |
| TEMP2 | 1662 | 605 | 130 | 1.5199 ± 0.8462 | 1.9103 ± 1.0353 | 137 | 85 |
| MTR2 | 1192 | 1801 | 130 | 1.9613 ± 0.8337 | 2.0487 ± 1.0160 | 117 | 310 |
| VIS3 | 206 | 356 | 130 | 1.6937 ± 1.1964 | 2.4646 ± 1.6593 | 23 | 115 |
| ATN | 1404 | 218 | 130 | 2.3340 ± 1.5716 | 1.7044 ± 0.9724 | 446 | 24 |
| TRI | 1285 | 1145 | 130 | 2.6817 ± 1.0842 | 2.1486 ± 0.9642 | 589 | 225 |
| FRON | 1507 | 1379 | 130 | 1.9833 ± 0.9875 | 1.8629 ± 0.8358 | 195 | 119 |

| | | | | | | | |
|---|---|---|---|---|---|---|---|
| aDM | 2279 | 871 | 130 | 1.6739 ± 0.7842 | 1.4732 ± 0.6637 | 120 | 17 |
| pDM | 386 | 295 | 130 | 2.1205 ± 1.0371 | 1.7691 ± 0.9698 | 86 | 29 |
| VIS4 | 6 | - | 130 | 3.1397 ± 0.7795 | - | 4 | - |
| lFP | 0 | - | 130 | - | - | 0 | - |
| rFP | 243 | - | 130 | 2.2077 ± 1.2804 | - | 74 | - |
| PREF | 835 | - | 130 | 1.7501 ± 1.1249 | - | 128 | - |
| SAL | 621 | - | 130 | 2.2390 ± 1.0813 | - | 141 | - |
| lPCG | - | 1357 | 130 | - | 1.7284 ± 0.9542 | - | 125 |
| lTPO | - | 723 | 130 | - | 1.6468 ± 1.0058 | - | 39 |
| Total | 19011 | 16223 | - | - | - | 2701 | 2169 |

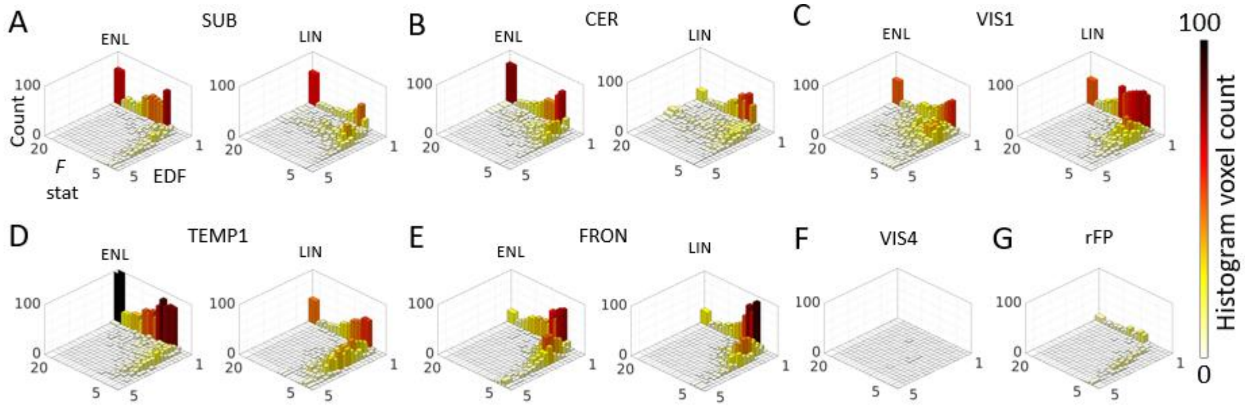


**Fig. S1.** Geometric complexity of explicitly nonlinear (ENL) and linear (LIN) intrinsic connectivity network developmental trajectories. (A-G) Two-dimensional histograms depict age association *F* statistics against effective degrees of freedom (EDF, quantifies nonlinearity of developmental trajectory) for significant ENL and LIN voxels for a subset of networks. Hue and bar height represent voxel count. SUB: subcortical. CER: cerebellum. VIS1: primary visual. TEMP1: primary temporal. FRON: frontal. VIS4: quaternary visual (ENL). rFP: right frontoparietal (ENL).

**Table S4.** Results of generalized additive model effective degrees of freedom (EDF, quantifies nonlinearity of developmental trajectory) comparisons between significant explicitly nonlinear (ENL) and linear (LIN) intrinsic connectivity network (ICN) voxels. Statistical tests were two-sided independent samples permutation tests with 5000 random permutations. Effect size is reported as Hedges's *g*. SUB: subcortical. CER: cerebellum. VIS1: primary visual. TEMP1: primary temporal. MTR1: primary sensorimotor. VIS2: secondary visual. TEMP2: secondary temporal. MTR2: secondary sensorimotor. VIS3: tertiary visual. ATN: dorsal attention. TRI: triangularis. FRON: frontal. aDM: anterior default mode. pDM: posterior default mode.

| **ICN** | ***n* (significant ENL voxels)** | ***n* (significant LIN voxels)** | **EDF difference (ENL – LIN)** | ***p*** | **Hedges's *g*** |
|---|---:|---:|---|---|---|
| SUB | 835 | 1014 | –0.3831 | < .001 | 0.4222 |
| CER | 1239 | 1139 | –0.3439 | < .001 | 0.4175 |
| VIS1 | 1748 | 1644 | 0.1931 | < .001 | 0.2195 |
| TEMP1 | 1606 | 1634 | –0.7683 | < .001 | 0.7608 |
| MTR1 | 1941 | 480 | 0.4037 | < .001 | 0.5401 |
| VIS2 | 16 | 1562 | 2.0974 | < .001 | 2.5359 |
| TEMP2 | 1662 | 605 | –0.3903 | < .001 | 0.4334 |
| MTR2 | 1192 | 1801 | −0.0874 | .0154 | - |
| VIS3 | 206 | 356 | –0.7709 | < .001 | 0.5117 |
| ATN | 1404 | 218 | 0.6296 | < .001 | 0.4182 |
| TRI | 1285 | 1145 | 0.5331 | < .001 | 0.5179 |
| FRON | 1507 | 1379 | 0.1204 | < .001 | 0.1312 |
| aDM | 2279 | 871 | 0.2007 | < .001 | 0.2666 |
| pDM | 386 | 295 | 0.3515 | < .001 | 0.3485 |
| Total | 19011 | 16223 | −0.0028 | .8078 | - |

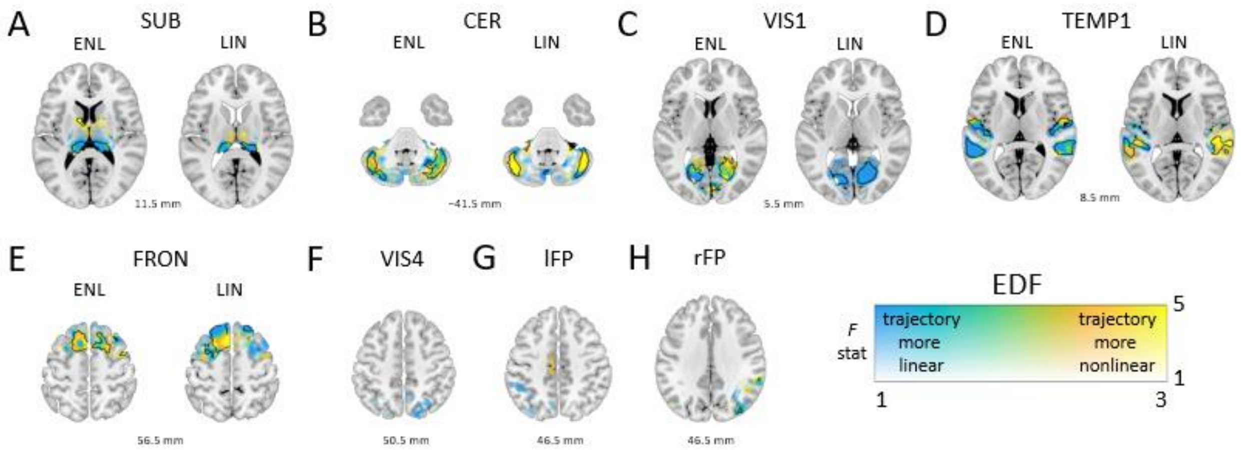


**Fig. S2.** Localization of explicitly nonlinear (ENL) and linear (LIN) intrinsic connectivity network associations with corrected postnatal age. (A-H) Spatial maps of voxel associations with age are plotted according to a dual-coded colormap for a subset of ENL and LIN networks, with transparency reflecting *F* statistic magnitude, hue representing model effective degrees of freedom (EDF; quantifies nonlinearity of developmental trajectory), and black contours indicating FDR-corrected significance ($q < .05$). Warmer hues indicate more complex, nonlinear developmental trajectories while cooler hues indicate less complex, approximately linear trajectories. Results are overlaid on the Montreal Neurological Institute 152 template with z coordinates listed relative to the origin. SUB: subcortical. CER: cerebellum. VIS1: primary visual. TEMP1: primary temporal. FRON: frontal. VIS4: quaternary visual (ENL). lFP: left frontoparietal (ENL). rFP: right frontoparietal (ENL).

**Table S5.** Results of comparisons between explicitly nonlinear (ENL) and linear (LIN) intrinsic connectivity network (ICN) statistical sensitivity to age. A two-sided McNemar's test was used to assess the overall ENL vs. LIN difference in statistical sensitivity (for all voxels tested across concordant ICNs), and differences in statistical sensitivity for concordant ICN pairs were assessed separately using two-sided McNemar's tests. Odds ratio (OR) is used as an indicator of effect size. SUB: subcortical. CER: cerebellum. VIS1: primary visual. TEMP1: primary temporal. MTR1: primary sensorimotor. VIS2: secondary visual. TEMP2: secondary temporal. MTR2: secondary sensorimotor. VIS3: tertiary visual. ATN: dorsal attention. TRI: triangularis. FRON: frontal. aDM: anterior default mode. pDM: posterior default mode.

| **ICN** | **No. significant voxels identified for ENL but not LIN** | **No. significant voxels identified for LIN but not ENL** | ***df*** | $\chi^2$ | ***p*** | **OR** |
|---|---|---|---|---|---|---|
| SUB | 280 | 492 | 1 | 57.6697 | < .00001 | 1.76 |
| CER | 461 | 250 | 1 | 62.0253 | < .00001 | 1.84 |
| VIS1 | 524 | 555 | 1 | 0.8341 | .3611 | - |
| TEMP1 | 560 | 555 | 1 | 0.0143 | .9046 | - |
| MTR1 | 1397 | 143 | 1 | 1019.5 | < .00001 | 9.77 |
| VIS2 | 1 | 1380 | 1 | 1375 | < .00001 | 1380 |
| TEMP2 | 959 | 112 | 1 | 668.2689 | < .00001 | 8.56 |
| MTR2 | 111 | 467 | 1 | 218.0363 | < .00001 | 4.21 |
| VIS3 | 49 | 238 | 1 | 123.1498 | < .00001 | 4.86 |
| ATN | 1199 | 37 | 1 | 1090.6 | < .00001 | 32.41 |
| TRI | 469 | 477 | 1 | 0.0518 | .82 | - |
| FRON | 597 | 654 | 1 | 2.5068 | .1134 | - |
| aDM | 929 | 229 | 1 | 421.9352 | < .00001 | 4.06 |
| pDM | 297 | 60 | 1 | 156.0112 | < .00001 | 4.95 |
| Total | 7833 | 5649 | 1 | 353.4705 | < .00001 | 1.39 |
